# Base edited “universal” donor CAR T cell strategies for acute myeloid leukaemia

**DOI:** 10.1101/2024.12.28.630573

**Authors:** Renuka Kadirkamanathan, Christos Georgiadis, Arnold Kloos, Annie Etuk, Roland Preece, Oliver Gough, Axel Schambach, Martin Sauer, Michael Heuser, Waseem Qasim

## Abstract

Acute myeloid leukaemia (AML) is often aggressive and life-threatening with limited curative options. Immunotherapies including chimeric antigen receptor (CAR) T cell approaches are under investigation, but high levels of disease heterogeneity remain a major hurdle to achieving durable responses. Targeting of multiple surface antigens may ensure complete immunological coverage of leukemic blast populations, but such antigens are often also present on healthy haematopoietic populations. To address likely aplasia, strategies can be designed to bridge CAR T cell therapies to allogeneic stem cell transplantation (allo-SCT), as demonstrated in recent anti-CD7 CAR T cell studies. Here we report that monotherapy using base edited (BE) ‘universal’ donor CAR T cells against CD33, CLL-1, or CD7 delivered inhibition of AML in immunodeficient mice when antigen expression was homogenous, but combined use of BE-CAR33 and BE-CARCLL-1 T cells was required to address heterogenous CLL-1^-/+^CD33^-/+^ disease. We also demonstrate that removal of shared CD7 antigens enabled compatibility of BE-CAR33 with BE-CAR7 T cells, including in a patient-derived xenograft (PDX) model of AML. Therapeutic strategies envisage ‘pick and mix’ applications of base edited universal CAR T cells in combinations determined by patient-specific antigen profiles. Such approaches also offer the possibility of deep, cell-based, de-bulking and preparative conditioning ahead of allo-SCT followed by donor-derived reconstitution.

## Introduction

Acute myeloid leukaemia (AML) is often an aggressive, life-threatening disease with limited curative treatment options [1]. Newly diagnosed patients typically receive multiple rounds of chemotherapy and those with “high-risk” disease or elevated minimal residual disease (MRD) may undergo allogeneic haematopoietic stem cell transplant (allo-SCT) with the aim of harnessing graft-versus-leukaemia (GvL) effects [2]. Approaches using chimeric antigen receptor (CAR) T cells are also under investigation, but their development and application is more challenging than for B-cell acute lymphoblastic leukaemia (B-ALL), especially in the context of disease heterogeneity and reduced immune cell fitness. Targetable antigens include CD33, CLL-1, CD123, CD7, CD38 and FLT-3, but expression of these on healthy tissues as well as disease populations suggests immunotherapy has to be time-limited to allow for eventual autologous recovery or donor derived reconstitution following haematopoietic SCT [3–12]. As an alternative to autologous CAR T cells, effectors can be generated from donors through an “off-the-shelf” approach, and use without HLA matching may be feasible following appropriate genome editing steps. We previously reported the use of cytidine base editing to develop ‘universal’ donor CAR T cells (BE-CAR7) against CD7, a molecule expressed on T-cell acute lymphoblastic leukaemia (T-ALL) and present on a large proportion of healthy lymphocytes. Multiplexed knockouts prevented expression of αβ T cell receptor (TCRαβ) to avoid graft-versus-host disease (GvHD), disrupted *CD7* to evade self-targeting through the anti-CD7 CAR, and removed *CD52* to confer resistance to the lymphodepleting antibody Alemtuzumab [13]. Similar anti-CD7 CAR T cell products are being investigated for their ability to mediate lymphodepletion and myeloablation ahead of allogeneic (allo-) SCT, and have been utilised to treat AML exhibiting CD7 antigen expression [14]. However, the majority of AML blasts have heterogenous antigen expression, necessitating the targeting of multiple antigens simultaneously [15]. Combinational approaches against B-cell malignancies have investigated various strategies ranging from the expression of different CARs via the transduction of T cells using multiple vectors, as well as bi-cistronic constructs or tandem configurations [16,17]. Our ‘off-the-shelf’ approach envisages the co-infusion of cells from pre-generated CAR T-cell banks with different target specificities. We therefore manufactured universal BE-CAR formulations against CD33 (siglec-3) or C-type lectin-like molecule-1 (CLL-1/CLEC12A), both of which are commonly over-expressed in childhood and adult AML [3,4]. Multiplexed base-editing of *TRBC*, *CD7* and *CD52* generated universal BE-CAR33 and BE-CARCLL-1 CAR T cells compatible with our existing BE-CAR7 products. Each BE-CAR product was investigated alone, or in combination, against heterogenous cell lines and primary AML, as a cell-based universal CAR T cell immunotherapy approach against AML.

## Methods

### Cell lines and primary cells

HEK293Ts, Kasumi-3, HL-60, U-937 and Molm-14 cell lines were obtained from ATCC and cultured according to the supplier’s instructions. Healthy donor peripheral blood collections were approved by University College London (UCL). Patient-derived xenograft (PDX) AML models were established and maintained at Hannover Medical School (MHH), Germany.

### Flow cytometry

When appropriate, AML cell lines and samples from *in vivo* studies were blocked with Human TruStain FcX (BioLegend, San Diego, CA, USA) prior to staining. Samples were acquired using the CytoFLEX cytometer (Beckman Coulter, High Wycombe, UK), CyAn^TM^ ADP analyser (Beckman Coulter), BD^®^ FACSymphony^TM^ cytometer (BD, Franklin Lakes, NJ, USA) or the BD^®^ LSR II flow cytometer. Data was analysed using FlowJo^TM^ software v10 (TreeStar Inc, Ashland, OR, USA).

### Preparation of 3^rd^ generation BE-CAR lentiviral vectors

3^rd^ generation lentiviral vector encoding BE-CAR7 was prepared as described previously [13]. Antigen binding regions of CAR33 and CARCLL-1 constructs comprised codon-optimised scFv sequences from variable-heavy and variable-light chain regions of the My96 anti-CD33 antibody clone (DrugBank ID DB00056) and the M26 anti-CLL-1 clone (US 2013/0295118) respectively, with a (GGGGS)_3_ linker. Each scFv was included upstream of the hinge and transmembrane domains of CD8_α_ and intracellular signalling domains of 4-1BB and CD3ζ. pTTB-CAR33 and pTTB-CARCLL-1 CRISPR-CAR self-inactivating lentiviral vectors included an internal PGK promoter, central polypurine tract (cppt) elements and mutated woodchuck hepatitis virus post-transcriptional regulatory element (WPRE). pTTB vectors incorporated a sgRNA cassette against *TRBC* embedded in the 3’ long terminal repeat (LTR) as previously described and were adapted for compatibility with base editing [18]. Self-inactivating (SIN) lentiviral vector stocks were produced via transient transfection of HEK293T cells using 3^rd^ generation packaging plasmids (Plasmid factory, Bielefeld, Germany) and a pseudotyped vesicular stomatitis virus G protein (VSV G) envelope plasmid. Supernatant was collected 48 hours later, filtered at 0.45 μm and concentrated through ultracentrifugation.

### SpCas9- and BE3-mediated genome editing

Synthetic sgRNAs were manufactured by Synthego (Redwood City, CA, USA) against *TRBC* (exon 1 - tryptophan to pmSTOP) 5’ CCCACCAGCTCAGCTCCACG 3’, *CD7* (exon 2 - glutamine to pmSTOP) 5’ CACCTGCCAGGCCATCACGG 3’ and *CD52* (exon 1 - splice site disruption) 5’ GGTTATGGTACAGGTAAGAG 3’, *CD33* (exon 1 - pmSTOP) 5’ TGACAACCAGGAGAAGATCG 3’ and *CLEC12A* (exon 1 – double stranded break formation) 5’ TGCTGGACGCCATACATGAG 3’. mRNA encoding codon optimised SpCas9 or codon optimised BE3 (coBE3) was supplied by TriLink BioTechnologies (San Diego, CA, USA) and BioNTech (Mainz, Germany) respectively.

### Generation of BE-CAR T cell effectors

MNCs were isolated from peripheral blood through Ficoll separation, activated with TransAct reagent (Miltenyi Biotec, Bergisch Gladbach, Germany) and cultured at a density of 1×10^6^ per mL in TexMACS (Miltenyi Biotec) media with 3% heat-inactivated human serum (Seralab, Sussex, UK) and 20 ng/mL human recombinant IL-2 (Miltenyi Biotec). BE-CAR7 products and CART19 control cells were made as previously described using pCCL-CAR7 or pTT-CAR19 vector, respectively, as described previously [13,19].

For BE-CAR33 and BE-CARCLL-1 products, cells were transduced with pTTB-CAR33 or pTTB-CARCCL1 24 hours after activation at MOI of 5 and cultured for a further 72 hours. Editing sgRNAs (*CD7* and *CD52* 10 μg/mL) and coBE3 (50 μg/mL) were delivered via electroporation using a Lonza 4D-Nucleofector^TM^ (EW138 pulse code). Cells were recovered for 16 hours at 30°C and returned to 37°C in G-Rex chambers as per the manufacturer’s instructions for a further 7 days. Cells underwent a TCRαβ depletion step (anti-TCRαβ antibody REA652 clone, anti-biotin microbeads and LD column, all Miltenyi Biotec) with an overnight recovery followed by cryopreservation in 45% TexMACS media (Miltenyi Biotec), 45% heat-inactivated human serum (Seralab), 10% DMSO.

### Molecular quantification of on-target genome editing

Genomic DNA was extracted using a DNeasy Blood and Tissue Kit (QIAGEN, Hilden, Germany), and 400-800bp fragments including the protospacer sequence were amplified via PCR using the following primers: *TRBC* fwd 5’ AGGTCGCTGTGTTTGAGC 3’ and *TRBC* rev 5’ CTATCCTGGGTCCACTCGTC 3’, *CD7* fwd 5’ TAGGTGAGACCGCCCCTCC 3’ and *CD7* rev 5’ GGGACCCTGAGAAGTCGATGC 3’, *CD52* fwd 5’ CTACCAAGACAGCCACGAAGAT 3’ and *CD52* rev 5’ TGCTCTCAGGAGAGAAGGCTG 3’. PCR products were purified using a QIAQuick PCR purification kit (QIAGEN) and sent for Sanger sequencing (Eurofins Genomics, Luxembourg City, Luxembourg). Rates of on-target C>T conversion were calculated using EDITR software (https://moriaritylab.shinyapps.io/editr_v10/).

### Flow-cytometry based co-culture assays

BE-CAR33 T-cell targets were labelled with 1μM CFSE at 5×10^6^/mL for 10 minutes at 37°C and combined with unlabelled BE-CAR7 effectors at an E:T of 1:1 for 16 hours in media without IL-2. Samples were harvested, left unwashed, stained with anti-CD2 antibody and DAPI to aid in the discrimination of dead cells and analysed using a Beckman Coulter CytoFLEX cytometer.

### 51Cr *in vitro* cytotoxicity assessments

Target cell lines were loaded with Cr^51^ (PerkinElmer, Waltham, MA, USA) for 1 hour at 37°C. BE-CAR T cells or untransduced T cells were co-cultured with targets at a range of E:Ts from 20:1 to 0.156:1. Maximum and spontaneous control wells were treated with 10X Triton X-100 (Sigma Aldrich, St. Lois, MO, USA), or media, respectively. All conditions were incubated at 4 hours at 37°C. Supernatants were then harvested, mixed with OptiPhase HiSafe 3 scintillation fluid (PerkinElmer) and incubated for 16 hours at room temperature. Levels of Cr^51^ release were measured using a microplate scintillation counter (Wallac 1450 MicroBeta TriLux, Perkin Elmer) and specific target lysis was calculated as [(experimental release – spontaneous release)/(maximum release – spontaneous release) x 100%].

### Cytometric bead array to measure BE-CAR *in vitro* cytokine release

Target cell lines were co-cultured with BE-CAR T cells or untransduced T cells at an E:T of 1:1, each at 1×10^6^ per mL, for 16 hours at 37°C in 20% IMDM. Supernatants were harvested from experimental wells. Sample preparation and acquisition was performed using the Human Th1/2/17 CBA kit (BD) and a BD^®^ LSR II flow cytometer, respectively. Data was then analysed using BD FCAP Array software v3.

### *In vitro* BE-CAR T cell activity against PDX samples

AML cells were derived from a newly diagnosed patient with t(3;3) rearrangement, classified as adverse risk (Europena LeukemiaNet, ELN 2022). BE-CAR33, BE-CARCLL-1 and untransduced effector cells were labelled with 1μM CFSE prior to co-culture with PDX cells at an E:T of 10:1 for 16 hours. Samples were harvested, left unwashed and first blocked with Human TruStain FcX^TM^ solution followed by staining with counting cocktail that contained CountBright^TM^ beads, BD Horizon^TM^ viability stain and 1μL of CD45 BV752 antibody. Acquisition was performed using a BD FACSymphony^TM^ cytometer and the stopping gate was set to 1,000 beads.

### Generation of mixed target antigen variants

Editing reagents (sgRNA at 10 μg/mL, coBE3 at 50 μg/mL or SpCas9 at 100 μg/mL) were transiently delivered via electroporation using a Lonza 4D-Nucleofector^TM^ (EN138 pulse code). Cells were recovered at 30°C for 16 hours and returned to 37°C. Phenotypic knockout of CLL-1 and CD33 were confirmed through flow cytometry 5 days following electroporation. Cells were then enriched for the CLL-1^-^CD33^+^ or CD33^-^CLL-1^+^ phenotype through fluorescence activated cell sorting (FACS) using the BD FACSAria^TM^ cell sorter.

### Generation of GFP expressing cell lines

Cells were transduced with 3^rd^ generation SIN pCCL lentiviral vector encoding a GFP-P2A-luciferase sequence at an MOI of 1. Fluorescent protein expression was confirmed 24 hours later through flow cytometry and cells were cultured for a further 7 days post-transduction. Fluorescent protein-expressing populations were then enriched for through FACS using the FACSAria^TM^ cell sorter (BD).

### *In vivo* BE-CAR activity against AML cell lines

Six-week-old NOD/SCID/ψc^-/-^ (NSG) mice were obtained from The Jackson Laboratory (Bar Harbor, ME, USA) (Charles River strain 005557). On day 0, mice were administered with 1×10^6^ Kasumi-3 GFP^+^LUC^+^ targets or 1×10^6^ HL-60 GFP^+^LUC^+^ (homogenous CLL-1^+^CD33^+^ or heterogenous CD33^-^CLL-1^+^/CLL-1^-^CD33^+^) via intravenous tail-vein injection. Tumour engraftment was measured on day 3 through bioluminescent imaging (BLI) with an IVIS Lumina III (Revvity, Waltham, MA, USA) *in vivo* imaging system. The following day, mice were then administered with BE-CAR effector cells, either alone (2×10^6^ CAR^+^ cells) or in a combination (4×10^6^ CAR^+^ cells), through intravenous tail-vein injection. Tumour growth was measured until the end of the experiment, at which point mice were sacrificed and bone marrow samples were processed for phenotypic analysis through flow cytometry. All animal studies were approved by the UCL Biological Services Ethical Review Committee and licensed under the Animals (Scientific Procedures) Act 1986 (Home Office, London, United Kingdom).

### *In vivo* BE-CAR activity in a PDX model of AML

Eight-to-twelve-week-old NOD/SCID/ψc^-/-^ NSG mice were sub lethally irradiated at 2.5Gy on day -21 and subsequently engrafted with AML PDX cells via tail-vein injection on day -20. AML cells were derived from a newly diagnosed patient with t(3;3) rearrangement, classified as adverse risk (ELN 2022). Written informed patient consent was obtained in accordance with the Declaration of Helsinki, and the study was approved by the local ethics review committee (ethical vote 1187-2011). The PDX mouse experiment adhered to the guidelines for animal care and use established by Hannover Medical School and was conducted with approval from the Lower Saxony state office for consumer protection, Oldenburg, Germany. Mice were kept under pathogen-free conditions at the central animal laboratory of Hannover Medical School. Peripheral bleeds were performed to measure levels of engraftment from day -1. Mice then received CAR19, BE-CAR7 or BE-CAR33 effector cells individually or combined on day 0. Serial peripheral bleeds were then repeated to track disease progression. Two weeks post CAR T infusion, one mouse from each group was sacrificed and bone marrow samples were processed for phenotypic analysis through flow cytometry.

### Statistical analysis

Statistical significance was determined through a one-way ANOVA with a Tukey multiple comparison post-hoc test or a Mann-Whitney U test using GraphPad (La Jolla, CA, USA) Prism software V9.0. Error bars represent standard error of the mean.

## Results

### Generation of universal BE-CAR T cells

BE-CAR7 T cells were generated as previously reported using coBE3 and sgRNAs targeting *CD7*, *TRBC* and *CD52* ahead of lentiviral pCCL-CAR7 transduction (Fig. 1. Supplementary Fig. 1A), and resulted in 51.0±5.3% BE-CAR7^+^TCRαβ^-^ cells (n=4) (Figs. 1B, D). Universal BE-CAR33 and BE-CARCLL-1 T cells were generated by first transducing activated T cells with pTTB-CAR lentiviral vectors incorporating a *TRBC* specific sgRNA expression cassette. Editing machinery was operational only when combined with coBE3 mRNA delivery and additional sgRNA sequences targeting *CD7 and CD52* were also delivered via electroporation (Fig. 1A, Supplementary Fig. 1A). Coupled TCRαβ disruption and CAR expression ensured that following magnetic bead depletion of residual TCRαβ^+^ cells, products were highly enriched as CAR33^+^TCRαβ^-^ T cells (mean 92.4±2.1%, n=10) or CARCLL-1^+^TCRαβ^-^ T cells (mean 94.9±1.0%, n=11) (Figs. 1C, D). Residual TCRαβ^+^ expression was 1.1±0.5% (n=4), 2.0±0.8% (n=10) and 1.0±0.3% (n=11) for BE-CAR7, BE-CAR33 and BE-CARCLL-1 products respectively (Fig. 1D). Residual CD7 expression of 29.8±4.6% (n=10) and 32.4±4.5% (n=11) BE-CAR33 and BE-CARCLL-1 cells was documented at end-of-manufacture, compared to BE-CAR7 T-cells where self-enrichment resulted in <1% CD7^+^ cells (n=4) (Figs. 1B, C, E). Residual CD52 expression was present on 2.9±1.5% (n=4), 6.6±2.7% (n=4) and 9.0±4.8% (n=5) BE-CAR7, BE-CAR33 and BE-CARCLL-1 cells, respectively (Figs.1B, C, E). BE-CAR T-cell phenotypes at the end of production captured CD45RA and CD62L expression to broadly define naïve (T_N_), central memory (T_CM_), effector memory (T_EM_) and terminally differentiated (T_EMRA_) T-cell subsets (Supplementary Fig. 1C). Flow cytometry for PD-1, BTLA, TIM-3 and/or LAG-3 upregulation provided exhaustion marker profiles (Supplementary Fig. 1D).

**Fig. 1.**
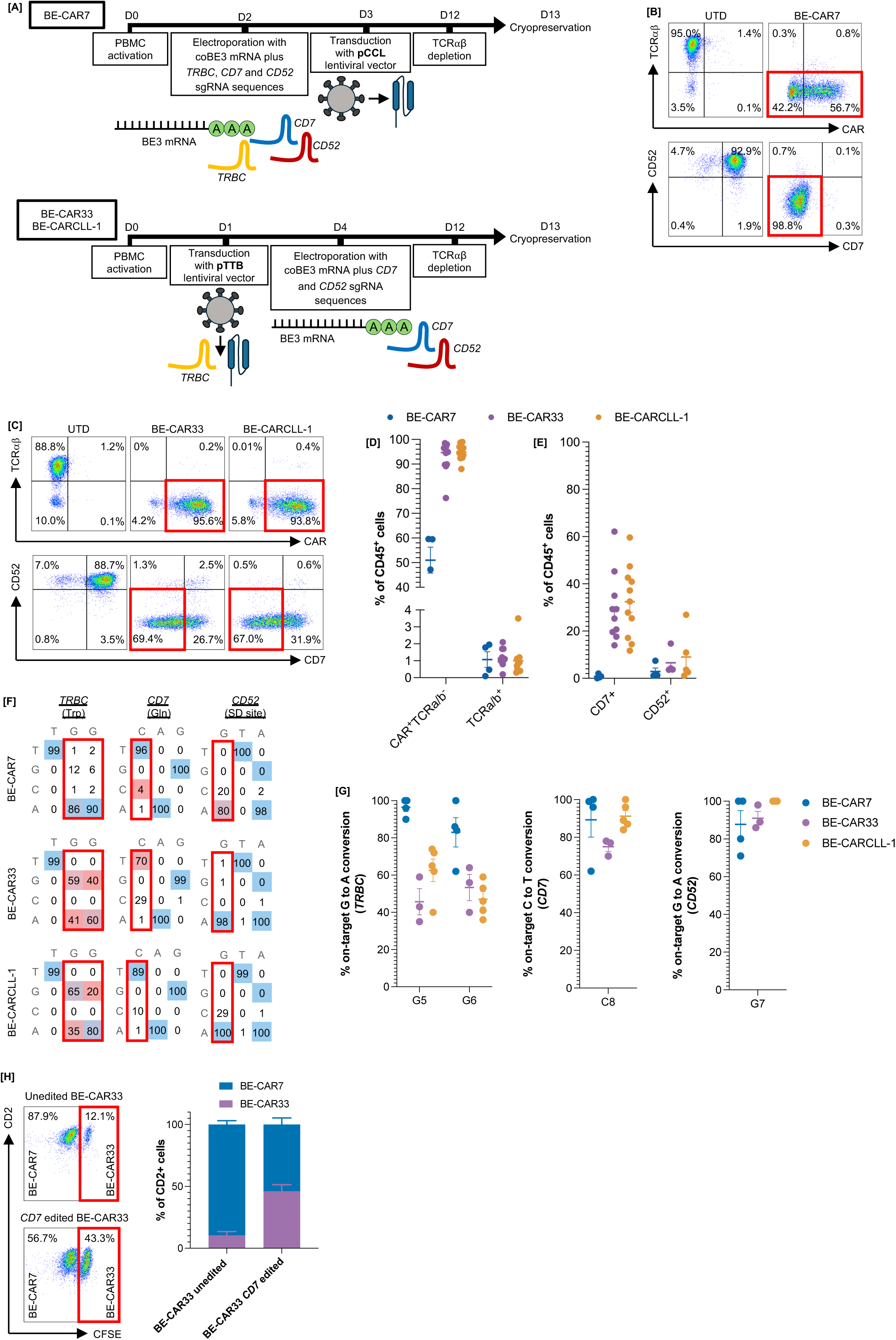
BE-CAR7, BE-CAR33 and BE-CARCLL-1 T cells to target AML. **[A]** For BE-CAR7 T cells, PBMCs were electroporated with codon optimised mRNA encoding the cytidine base editor BE3 and sgRNAs against *TRBC* (to introduce a premature stop codon), *CD7* (to introduce a premature stop codon) and *CD52* (to disrupt a splice donor site). Cells were then transduced at an MOI of 5 with a pCCL lentiviral vector encoding anti-CD7 CAR and residual TCRɑβ^+^ cells were subsequently removed through magnetic bead and column depletion. For BE-CAR33 and BE-CARCLL-1 T cells, PBMCs were transduced at an MOI of 5 with a pTTB lentiviral vector encoding the anti-CD33 or anti-CLL-1 CAR alongside a *TRBC* specific sgRNA expression cassette, and were subsequently electroporated to transiently deliver co BE3 mRNA and sgRNAs against *CD7* and *CD52*. Residual TCRɑβ^+^ cells were subsequently removed through magnetic bead and column depletion. Created with BioRender.com. **[B]** CD45^+^ untransduced and end-of-manufacture BE-CAR7 T cells displaying cell surface CAR and/or TCRαβ (top row) and CD7 and/or CD52 (bottom row) expression following electroporation, transduction and magnetic column depletion steps. Cells displaying CAR^+^TCRαβ^-^ and CD7^-^CD52^-^ phenotypes are highlighted in red. **[C]** CD45^+^ untransduced and end-of-manufacture BE-CAR33 and BE-CARCLL-1 cells displaying cell surface CAR and/or TCRαβ (top row) and CD7 and/or CD52 (bottom row) expression following transduction with pTTB-CAR, electroporation and magnetic bead depletion. Cells displaying CAR^+^TCRαβ^-^ and CD7^-^CD52^-^ phenotypes are highlighted in red. **[D]** End-of-manufacture BE-CAR7, BE-CAR33 and BE-CARCLL-1 T cells displaying a CAR^+^ TCRαβ^-^ phenotype and residual TCRαβ^-^ expression, measured through flow cytometry. **[E]** End-of-manufacture BE-CAR7, BE-CAR33 and BE-CARCLL-1 T cells displaying cell surface CD7 and CD52 expression, measured through flow cytometry. **[F]** EditR plots demonstrating the percentage of on-target base conversion at *TRBC*, *CD7* and *CD52* editing sites. **[G]** On-target base conversion at *TRBC*, *CD7* and *CD52* editing sites determined through EditR analysis of Sanger sequencing results. **[H]** CFSE labelled BE-CAR33 targets (T), unedited or edited to disrupt CD7 expression, and BE-CAR7 effectors (E) remaining after co-culture for 16 hours at an E:T of 1:1.

Molecular mapping performed by Sanger sequencing revealed G_5_>A base conversions for *TRBC* of 96.5±2.4% for BE-CAR7 (n=4)), 45.7±7.1% (n=3) for BE-CAR33 and 62.6±6.1% (n=4) for BE-CARCLL-1 conversion (Figs. 1F, G). At the *CD7* site C_8_>T conversions were quantified as 89.3±9.1% for BE-CAR7 (n=4), 75.0±2.5% for BE-CAR33(n=3) and 91.2±3.0% (n=5) for BE-CARCLL-1 and G_7_>A conversions at the *CD52* site were 87.8±7.3% for BE-CAR7 (n=4), 91.0±3.6% for BE-CAR33 (n=3) and 100±0% (n=3) for BE-CARCLL-1 T cells (Figs. 1F, G).

Protective effects of *CD7* disruption on BE-CAR33 products against BE-CAR7-mediated fratricide were confirmed in a 16-hour co-culture assay where residual CD7^+^CAR33^+^ cells were targeted by BE-CAR7 cells, whereas CD7^-^CAR33^+^ cells largely evaded cytotoxicity (Fig. 1H, Supplementary Fig. 1E).

### BE-CAR T cells against AML with homogenous antigen profiles

Functional studies were performed to assess *in vitro* cytotoxicity of BE-CAR7 T cells against Kasumi-3 cells, whilst BE-CAR33 and BE-CARCLL-1 T cells were tested against Molm-14, U-937 and HL-60 cells (Supplementary Figs. 2A-C). CD33 and CLL-1 antigen density was documented for each relevant cell line (Supplementary Figs. 2D) before ^51^Cr release assays (Fig. 2A) and cytokine release quantification (Fig. 2B) after co-culture with effectors across a range of target ratios. Robust CAR-mediated target cell lysis was documented for each BE-CAR product, and cytokine release profiles captured. In addition to cell lines, targeting of primary human AML expressing CLL-1^high^ and CD33^high^ by BE-CAR33 and BE-CARCLL-1 T cells was demonstrated in overnight co-culture experiments (Supplementary Fig. 2E).

**Fig. 2.**
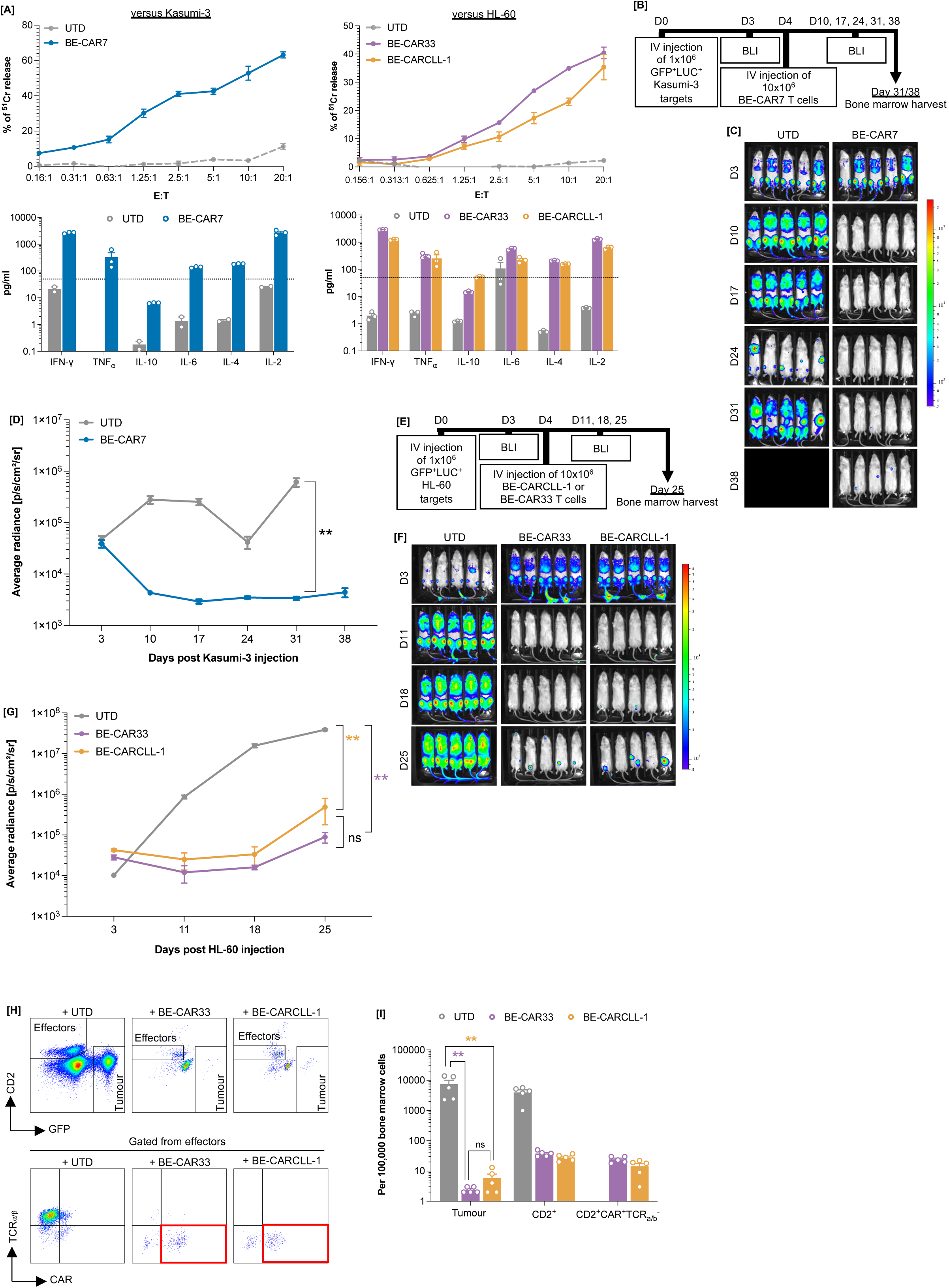
BE-CAR7, BE-CAR33 and BE-CARCLL-1 T cells exhibited specific anti-leukaemia effects *in vivo* against AML with homogenous antigen expression. **[A]** (Top) Normalised ^51^Cr release from Kasumi-3 (left) or HL-60 (right) targets when co-cultured with BE-CAR7, and BE-CARCLL-1 or BE-CAR33 T cells for 4 hours at E:Ts ranging from 20:1 to 0.156:1. Untransduced effector controls were included to exclude non-specific lysis. (Bottom) Th1- and Th2-cytokines released from untransduced, BE-CAR7, BE-CARCLL-1 or BE-CAR33 T cells when co-cultured with Kasumi-3 (left) or HL-60 (right) at an E:T of 1:1 for 16 hours. Cytokine release below 50 pg/mL was considered background (dotted line). **[B]** A humanised xenograft murine model of AML with a GFP^+^LUC^+^ Kasumi-3 cell line was used to assess BE-CAR7 anti-leukaemia effects *in vivo*. **[C]** Bioluminescent signals from mice that received 1×10^6^ Kasumi-3 targets on day 0, and 10×10^6^ effectors on day 4. **[D]** Average radiance values (p/s/cm^2^/sr) of each animal following injection with Kasumi-3 targets, measured through BLI. Area under the curve was calculated for each experimental group and values compared using a one-way ANOVA with Tukey multiple comparison post-hoc. ** p < 0.01. **[E]** A humanised xenograft murine model of AML with a GFP^+^LUC^+^ HL-60 cell line was used to assess BE-CAR33 and BE-CARCLL-1 anti-leukaemia effects *in vivo*. **[F]** Bioluminescent signals from mice that received 1×10^6^ HL-60 targets on day 0, and 10×10^6^ effectors on day 4. **[G]** Average radiance values (p/s/cm^2^/sr) of each animal following injection with HL-60 targets, measured through BLI. Area under the curve was calculated for each experimental group and values compared using a one-way ANOVA with Tukey multiple comparison post-hoc. ** p < 0.01, ns; nonsignificant. **[H]** (Top) CD45^+^GFP^+^ tumour and CD2^+^ effectors detected in bone marrows of mice from each group. (Bottom) CD45^+^ bone marrow CD2^+^ effectors demonstrating cell surface CAR and/or TCRɑαβ expression detected in bone marrows of mice from each group. **[I]** Normalised numbers of CD45^+^GFP^+^ tumour and CD2^+^/CD2^+^CAR^+^TCRαβ-effector cells in the bone marrow of mice from each group. ** p < 0.01, ns; non-significant. Values were compared using a Mann-Whitney U test.

Next, *in vivo* anti-leukaemia effects of each BE-CAR were assessed in a humanised xenograft murine model of AML. For BE-CAR7 T cells, 6-week-old NSG mice received 1.0×10^6^ GFP^+^LUC^+^ Kasumi-3 cells (n=10) via tail-vein injection on day 0, and BLI was utilised to confirm disease engraftment on day 3. AML-bearing mice went onto receive 10×10^6^ BE-CAR7 T cells (n=5) or untransduced cells (n=5) on day 4 with serial BLI taking place on days 10, 17, 24, 31 and 38 to monitor disease progression (Figs. 2B, C). Potent leukaemia clearance was observed in mice treated with BE-CAR7 T cells when compared with untreated mice (p < 0.0001) (Fig. 2D). Separately, for BE-CAR33 and BE-CARCLL-1 T cells, 6-week-old NGS mice received 1.0×10^6^ GFP^+^LUC^+^ HL-60 cells (n=15) via tail-vein injection on day 0, and BLI was utilised to confirm disease engraftment on day 3. AML-bearing mice went on to receive 10×10^6^ BE-CAR33 (n=5) or BE-CARCLL-1 (n=5) T cells, or untransduced cells (n=5) on day 4 with serial BLI taking place on days 11, 18 and 25 to monitor disease progression (Figs. 2E, F). Leukaemia progression was again inhibited in mice treated with either BE-CAR when compared with untreated mice (p<0.01), with low numbers of GFP^+^LUC^+^ HL-60 cells detected in the bone marrows of treated mice when compared to the control group (p<0.01) (Figs. 2H, I). Persistence of BE-CAR33 and BE-CARCLL-1 T cells was also documented, with CD45^+^CD2^+^ T-cell effectors maintaining their CAR^+^TCRαβ^-^ phenotype (Figs. 2H, I).

### BE-CAR33 and BE-CARCLL-1 T cells effects against heterogenous AML *in vivo*

Combined targeting of CD33 and CLL-1 may offer a valuable strategy against heterogenous disease as both markers are often upregulated on AML, although complete expression of either antigen may not be present across the entire blast population. We investigated use of BE-CAR33 alongside BE-CARCLL-1 T cells *in vivo* against xenografted human CD33^+/-^ CLL-1^+/-^ AML. Mice were engrafted with equal numbers of HL-60 variants, edited using CRISPR/Cas9 to express either CD33 (CD33^+^CLL-1^-^) or CLL-1 (CLL-1^+^CD33^-^) (Supplementary Figs. 3A, B). BLI confirmed tumour engraftment on day 3 and AML-bearing mice went on to receive tail-vein injections of untransduced control T cells (n=5) or effector cells comprising BE-CARCLL-1 (n=5) or BE-CAR33 T cells (n=5) alone, or combined at equal numbers (n=5) (Fig. 4A). Mice receiving BE-CAR33 T cells as a monotherapy and untreated controls exhibited significant disease progression by day 25 when compared with animals receiving combined BE-CAR (p<0.05, p<0.005 respectively) (Fig. 4C). Mice sacrificed on day 25 due to disease progression had high levels of bone marrow disease detectable through flow cytometry. Mice receiving BE-CAR33 or BE-CARCLL-1 T cells as a monotherapy were able to clear cells expressing their corresponding target antigen with remaining disease mostly demonstrating CD33^-^CLL-1^+^ or CLL-1^-^CD33^+^ phenotypes, respectively. In contrast, an average radiance value of 1.2×10^7^ p/s/cm^2^/r was set as a threshold for tumour progression in mice treated with combined BE-CAR T cells, and 4 out of 5 reached this by day 39. Potent disease clearance was observed in the bone marrow of mice treated with combined BE-CAR T cells when compared with BE-CAR33 (p<0.01) or BE-CARCLL-1 (p<0.01) monotherapy groups. Importantly, very few residual HL-60 cells displaying loss of both CAR target antigens were detected in the bone marrows of mice from all combined BE-CAR T-cell groups (Fig. 4G). These findings together demonstrate combined anti-leukaemia effects of BE-CARCLL-1 and BE-CAR33 T cells can inhibit heterogenous AML more effectively than monotherapy strategies.

**Fig. 3.**
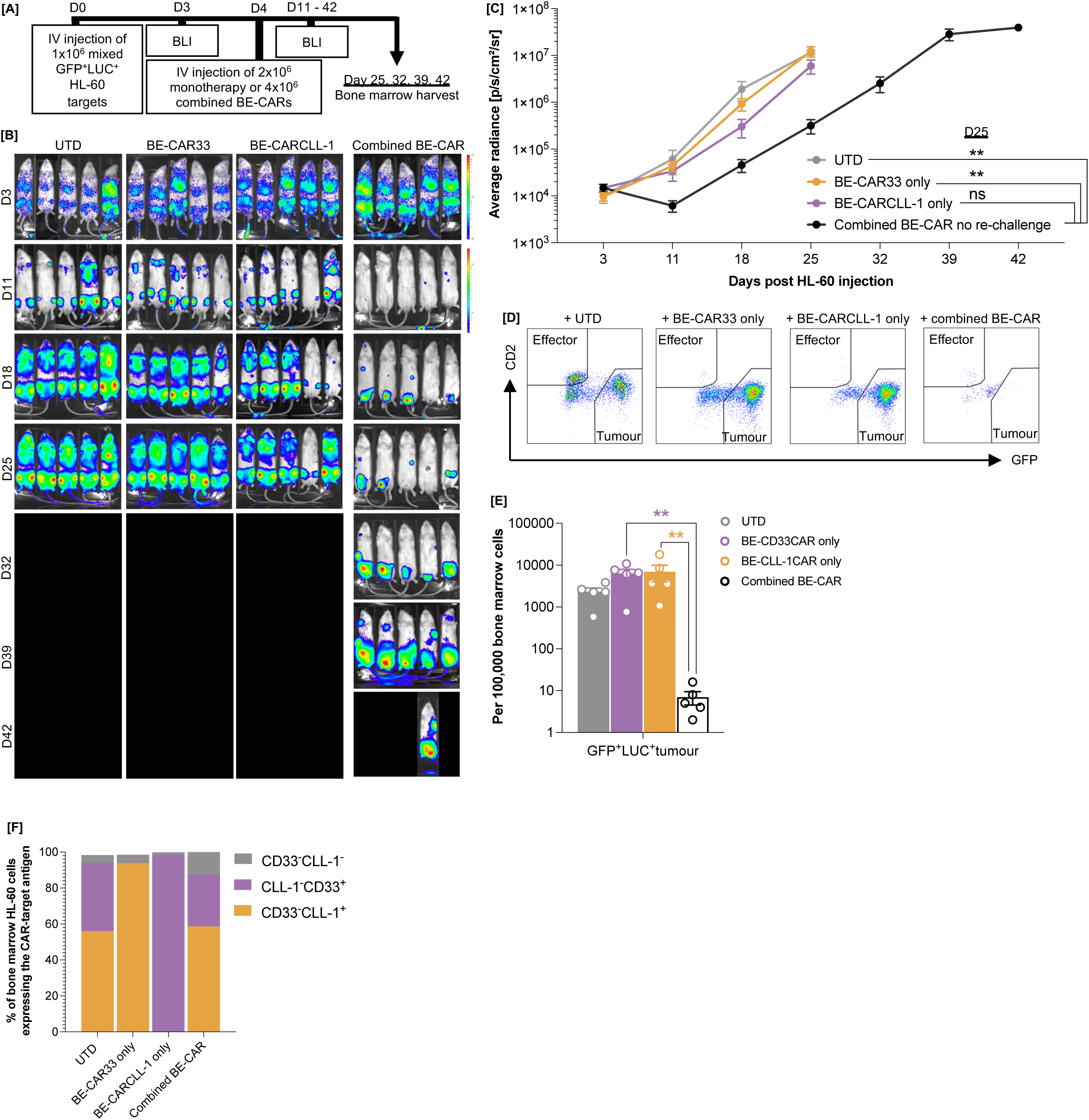
BE-CAR33 and BE-CARCLL-1 effectors exert combinatorial effects to clear heterogenous AML *in vivo*. **[A]** A humanised xenograft model of heterogenous AML with GFP^+^LUC^+^ variants of CRISPR-engineered CLL-1^-^CD33^+^ and CD33^-^CLL-1^+^ HL-60 cell lines was used to assess a combined dosing strategy with BE-CAR33 and BE-CARCLL-1 effectors. **[B]** Bioluminescent signals from mice that received 1×10^6^ CD33^-/+^CLL-1^-/+^HL-60 targets on day 0, and 10×10^6^ effectors on day 4. **[C]** Average radiance values (p/s/cm^2^/sr) of mice injected with 2×10^6^ monotherapy (n=5) or 4×10^6^ combined (n=5) BE-CARCLL-1 and/or BE-CAR33 effectors. ** p < 0.01, ns; non-significant. Values were compared using a one-way ANOVA with Tukey multiple comparison post-hoc. **[D]** CD45^+^GFP^+^ tumour cells detected in bone marrows of mice from each treatment group. **[E]** Normalised CD45^+^GFP^+^ tumour and CD2^+^ effector cell counts detected in bone marrow of mice from each treatment group. ** p < 0.01. Values were compared using a Mann-Whitney U test. **[F]** Percentage of CD45^+^GFP^+^ tumour detected in the bone marrow of mice from each treatment group demonstrating a CLL-1^-^CD33^+^, CD33^-^CLL-1^+^ or CD33^-^CLL-1^-^ phenotype.

**Fig. 4.**
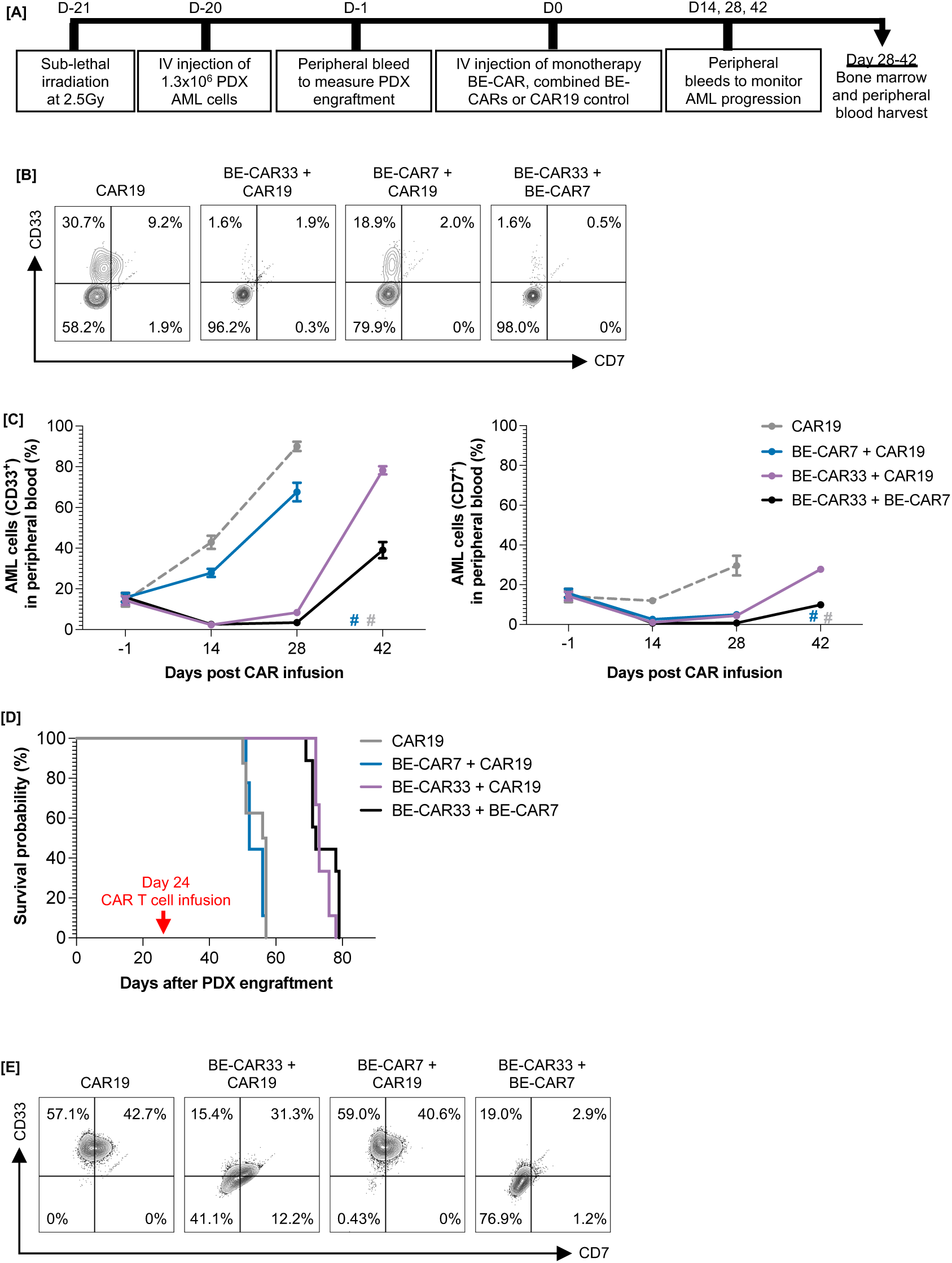
Removal of CD7 enables compatibility of BE-CAR7 and BE-CAR33 in a PDX model of AML. **[A]** A PDX model of CD33^high^CD7^low^ AML. Mice were sub lethally irradiated on day-21 prior to engraftment with 1×10^6^ PDX cells. Peripheral bleeds were performed to monitor PDX engraftment and disease progression through flow cytometry. **[B]** PDX cells located in peripheral blood of mice from each treatment group displaying CD33^-/+^CD7^-/+^ expression. **[C]** CD33^+^ (left) and CD7^+^ (right) PDX cells detected in the peripheral blood of mice from each treatment group. **[D]** Kaplan-Meir survival plot of mice treated with CAR19 control T cells, BE-CAR7 or BE-CAR33 cells, both in isolation and combination, on day 24 post engraftment with PDX cells. **[E]** Percentage of PDX cells detected in the bone marrows of mice from each treatment group demonstrating CD33^+^CD7^-^ or CD33^+^CD7^+^ disease phenotypes.

### BE-CAR33 activity in a PDX model of AML following co-infusion with BE-CAR7

A combinatorial strategy was tested *in vivo* to examine the feasibility of using BE-CAR33 T cells in the presence of BE-CAR7 T cells, noting that the latter is already being applied in human studies against T-ALL, and can offer lymphodepleting effects as well as direct anti-leukaemia activity against CD7^+^ AML. For these experiments, BE-CAR33 T cells were base edited to remove CD7 to prevent targeting by BE-CAR7 T cells and both populations were also edited to remove *CD52* in case of co-infusion in the presence of alemtuzumab. We utilised a PDX murine model where CD33^high^ CD7^low^ primary AML cells were engrafted into Eight-twelve week-old NSG mice following sub-lethal (2.5Gy) irradiation (Fig. 4A). Peripheral blood sampling confirmed AML engraftment three weeks later and effectors were injected in the following groups: CAR19 control T cells alone (n=9) or in combination with either BE-CAR33 (n=10) or BE-CAR7 (n=10), and BE-CAR33 in combination with BE-CAR7 T cells (n=10). Serial peripheral bleeds were then performed on day 14, 28 and 42 post-effector infusion to track disease progression. By day 14, groups that had received BE-CAR33 T cells exhibited delayed disease progression compared to mice that received BE-CAR7 and/or control CAR19 cells (Fig. 4B). Importantly, BE-CAR33 effectors continued to exert their anti-leukaemia effects in the presence of BE-CAR7 in the co-infusion group, confirming successful evasion of fratricidal effects through removal of CD7 (Figs. 4C, D). Treatment with BE-CAR33 cells significantly prolonged survival of PDX mice (Fig. 4D), and bone marrow analysis confirmed clearance of the majority of CD33^+^ disease two weeks post co-infusion with BE-CAR33 and BE-CAR7 T cells. Residual blasts were noted to be CD33^dim^CD7^low^ in the bone marrow demonstrating activity of BE-33CAR product (Fig. 4E). This data together confirms the anti-leukaemia effects of CD7 edited BE-CAR33 T cells were preserved and functional in the presence of BE-CAR7 T cells.

## Discussion

Children and adults with relapsed or refractory AML continue to face dismal long term survival outcomes with current therapy regimens [1]. CAR T cells offer a promising alternative but targeting of a single antigen may not be as effective as with B- or T-cell malignancies due to high levels of inter- and intra-patient phenotypic heterogeneity [15]. A lack of leukaemia-specific antigens also remains problematic, and target antigen expression inevitably overlaps with healthy haematopoietic or other tissues; for example, autologous and allogeneic CAR T cell therapies targeting the interleukin-3 receptor (CD123) have been investigated in clinical trials, but with notable toxicities and variation across age groups [20–22]. Other antigens of interest may also be present on precursors or mature granulocytes, macrophages and T cells, and myeloablative and lymphodepleting CAR T cell effects can create protracted risks of cytopenia and opportunistic infections [23,24]. In addition, disease heterogenicity suggests that CAR T-cell therapies against multiple leukaemia antigens may be needed to ensure complete coverage of disease profiles of both childhood and adult AML. CAR T cells used in combination may address heterogenicity and mediate disease clearance ahead of donor-derived stem cell rescue.

CD33 is broadly expressed on AML blasts and leukaemia stem cells (LSCs) and has been targeted using the drug conjugated humanised anti-CD33 monoclonal antibody (mAb) Gemtuzumab ozogamicin (GO) [25,26]. A number of alternative anti-CD33 antibody drug conjugates (ADCs), radiolabelled antibodies, bi-specific antibodies and CAR T cells carrying GO-derived binders have also been investigated clinically [27–30]. Qin et al. encountered toxicities in their murine studies and instead proposed a Lintuzumab-derived CAR T product with a CD28-CD3ζ signalling domain, and results from a corresponding clinical study in children and young adults suggested manageable toxicities with complete responses and myeloid aplasia reported in 2 out of 6 patients at higher doses [31,32]. Our GO-derived BE-CAR includes 41BB-CD3ζ signalling domains and successfully inhibited AML in humanised mice without obvious toxicity, including when used in combination with other BE-CAR products. Taking into account extensive pre-existing clinical safety data from anti-CD33 antibody studies, BE-CAR33 has been adopted for early phase clinical investigation in an allogeneic off-the-shelf approach in children (NCT05942599). Clinical experience with similar autologous anti-CD33 CAR T cell products have noted difficulties with apheresis harvests in heavily pre-treated patients [33]. Experience is limited to small series reports; Tambaro et al. treated 3 patients, describing cytokine release syndrome (CRS) and immune effector cell-associated neurotoxicity syndrome (ICANS), whilst Wang et al. reported transient blast reduction in a single patient [34]. Rapidly produced anti-CD33 CAR T cells with membrane bound IL-15 (mbIL-15) and a suicide switch have also been tested in a phase 1 clinical trial with documented objective responses in AML patients who received cells following lymphodepletion [35]. Recently, Applebaum et al. reported variant CD33 binders with rapamycin-regulated dimerization for controlled expression, and clinical endpoints were consistent with T-cell expansion alongside signs of anti-leukaemia activity [36].

CLL-1 is a type II transmembrane C-type lectin-like receptor that is often aberrantly expressed on AML blasts, as well as on LSCs [3,37]. CLL-1 targeting CAR T cells have been documented in clinical case reports; Zhang et al. reported complete responses in a small number of children, including sustained morphologic remission following an allo-SCT [38–40]. Separately, six out of nine adults were reported to be in ongoing complete remission three months following treatment [41]. Our BE-CARCLL-1 product was designed using the anti-CLL-1 mAb clone M26 following promising reports from both studies. A separate CD28-CD3ζ CAR designed with the same anti-CLL-1 mAb clone is also under clinical investigation (NCT04219163) [42]. Targeting of CLL-1 alongside CD33 has also been investigated, including via a self-cleaving P2A peptide sequence allowing for a 1:1 expression of both CAR constructs [43]. Six of nine patients were reported as MRD negative following CAR T-cell treatment and allo-SCT.

We also investigated the use of anti-CD7 CAR T cells as part of a combinatorial approach. CD7 is an Ig-superfamily molecule that is highly expressed on T cells and NK cells and some AML cases, with several anti-CD7 CAR T cells under clinical investigation to treat CD7^+^ disease [44]. We previously reported the generation of fratricide- and lymphodepletion-resistant universal BE-CAR7 T cells, developed originally to treat CD7^+^ T-cell malignancies [13]. Multiple studies, including one at our centre, have reported profound lymphodepletion and bone marrow aplasia post CAR T-cell infusion [45]. There are also reports of successful haploidentical-SCT without the need for additional myeloablative pre-conditioning post anti-CD7 CAR therapy [14]. We anticipate BE-CAR7 may offer dual benefits of targeting CD7^+^ disease and augmenting lymphodepletion and bone marrow conditioning ahead of bridging allo-SCT.

For BE-CAR33 and BE-CARCLL-1 products, our CRISPR-CAR vector configuration carrying a sgRNA cassette embedded in the vector LTR, previously deployed to manufacture our TT52CAR19 product for B-ALL, was combined with cytidine base editing. This enabled *TRBC* knockout to be coupled to CAR expression allowing for a highly enriched CAR^+^TCRαβ^-^ product [18]. Delivery of additional uncoupled sgRNAs at the time of electroporation along with BE mRNA rendered the cells resistant to pre-conditioning serotherapy with anti-CD52 antibodies such as alemtuzumab and removed shared T-cell antigen CD7. Each BE-CAR product was able to clear respective antigen-expressing AML targets with high efficiency alongside concordant elevation of Th1 cytokines *in vitro*, with anti-leukaemia potency also corroborated *in vivo* against human xenograft murine models of AML. Previous combined target antigen targeting of Molm-13, HL-60 and KG1a AML cell lines noted improved survival following combined dosing with anti-CD33 and anti-CLL-1 CAR T cells compared to monotherapy options [42]. Targeted removal of CD33 and CLL-1 form HL-60 cells provided a heterogenous model to assess specific BE-CAR products *in vivo*. Monotherapy treatment groups cleared target positive disease but succumbed to antigen negative disease, whilst combined infusions of BE-CAR33 and BE-CARCLL-1 cells were able to control disease progression. Furthermore, in a primary AML model, CD33^high^CD7^low^ disease was controlled with longer overall survival in groups receiving BE-CAR33. These effectors persisted and remained operational in the presence of BE-CAR7 given the removal of CD7 from both products. Whilst our model could not assess BE-CAR7 effects against AML directly, the preservation of BE-CAR33 effects was important to establish following co-infusion of the two products. We envisage adoption of “universal” BE-CAR33, BE-CARCLL-1 and BE-CAR7 products for AML eradiation or control ahead of transplant. Deep preparative conditioning ahead of programmed transplant and donor-derived immunological reconstitution may offer valuable routes to treatment for heterogenous hard-to-treat disease.

Genome editing has unlocked opportunities for engineering of CAR T cells, and may be further exploited for modification of shared antigens on healthy HSCs to support persistence during anti-leukaemia therapy. CRISPR-mediated removal of CD33 on healthy HSCs is being investigated in combination with GO antibody therapy (NCT04849910), whilst base editing for disruption of FLT-3 and CD45 epitopes has been modelled to allow evasion of CAR-T effects [46–48]. However, in heavily treated AML patients, modification of autologous HSC may be challenging in subjects who may harbour underlying cancer predispositions, or who have been heavily pre-treated. In the first instance, allogeneic universal donor BE-CAR T cells, alone or in combination, offer innovative routes to disease clearance and remission if planned carefully ahead of allo-SCT and donor derived reconstitution.

## Acknowledgements

This work was funded by the Wellcome Trust (215619/Z/19/Z), Medical Research Council (MR/X004619/1) and MRC DPFS (MR/V03961X/1), National Institute of Health Research (NIHR) and Great Ormond Street Biomedical Research Centre (RP-2014-05-007), BRC (IS-BRC-1215-20012) and The Child Health Research CIO PhD Studentship 2020/2021 (182120). The views expressed are those of the author(s) and not necessarily those of the NHS, the NIHR or the Department of Health. This study was further supported by grants HE 5240/6-1, HE 5240/6-2 to MH from DFG, grant 16 R/2021 to MH from DJCLS, and grant 70114189, 70114478 and 70114706 to MH from Deutsche Krebshilfe. We would like to thank Ailsa Greppi and Kyle O’Sullivan for their invaluable technical support with *in vivo* studies, Dr. Ayad Eddaudi at UCL Joint Great Ormond Street Institute of Child Health, supported by the Great Ormond Street Children’s Charity (GOSHC). Anthony Nolan Trust (803716/ SC038827) supplied healthy blood cell donations.

## Author contributions

W.Q., C.G., A.S., M.S. and M.H. designed research; R.K., C.G., A.K., A.E., R.P., O.G., M.H., and W.Q. performed research and data analysis; R.K., C.G. and W.Q. wrote the manuscript. All authors have reviewed and approved the manuscript.

## Data availability

The datasets generated during and/or analysed during the current study are available from the corresponding author on reasonable request.

