## Supplementary Data for "Base edited “universal” donor CAR T cell strategies for acute myeloid leukaemia"

### Base edited "universal" donor CAR T cell strategies for acute myeloid leukaemia – supplementary methods and results

#### Supplementary methods

##### Flow cytometry

The following antibodies were utilised: CD45 (HI30 clone, Miltenyi, HI30/5B1 clone, BioLegend), CD8 (REA734 clone, Miltenyi), CD4 (M-T466 clone, Miltenyi, SK3 clone, BioLegend), CD2 (LT2 clone, Miltenyi, RPA-2.10 clone, BioLegend), $TCR_{\alpha/\beta}$ (BW242/412 clone and REA-652 clone, Miltenyi, IP26 clone, BioLegend), CD52 (HI186 clone, BioLegend), CD7 (CD7-6B7 clone, BioLegend), CD45RA (REA1047 clone, Miltenyi), CD62L (145/15 clone, Miltenyi), CCR7 (FR 11-11E8 clone, Miltenyi), PD-1 (EH12.2H7 clone, BioLegend), TIM-3 (7D3 clone, BioLegend), LAG-3 (T47-530 clone, BioLegend), BTLA (MIH26 clone, BioLegend, J168-540 clone, BD), CD69 (FN50 clone, BioLegend), CLL-1 (50C1 clone, BioLegend), CD33 (WM53 clone, BioLegend). To measure cell surface CAR expression, samples were stained with biotin-conjugated goat anti-mouse IgG $F(ab)_2$ fragment (Jackson Immunoresearch, Stratech Scientific Limited) followed with PE-conjugated streptavidin (Miltenyi). The following isotype controls were utilised: anti-mouse IgG1κ (MOPC-21 clone) and anti-mouse IgG2aκ (MOPC-137 clone), both from BioLegend.

##### SpCas9- and BE3-mediated genome editing

Synthetic sgRNAs, comprising a 20-nt protospacer sequence, 80-nt CRISPR scaffold and 2’O-methyl 3’ phosphorothioate modifications were manufactured by Synthego (California, US) by automated solid-phase synthesis and eluted into nuclease-free Tris-EDTA buffer.

#### Supplementary results

**Supplemental figure 1**

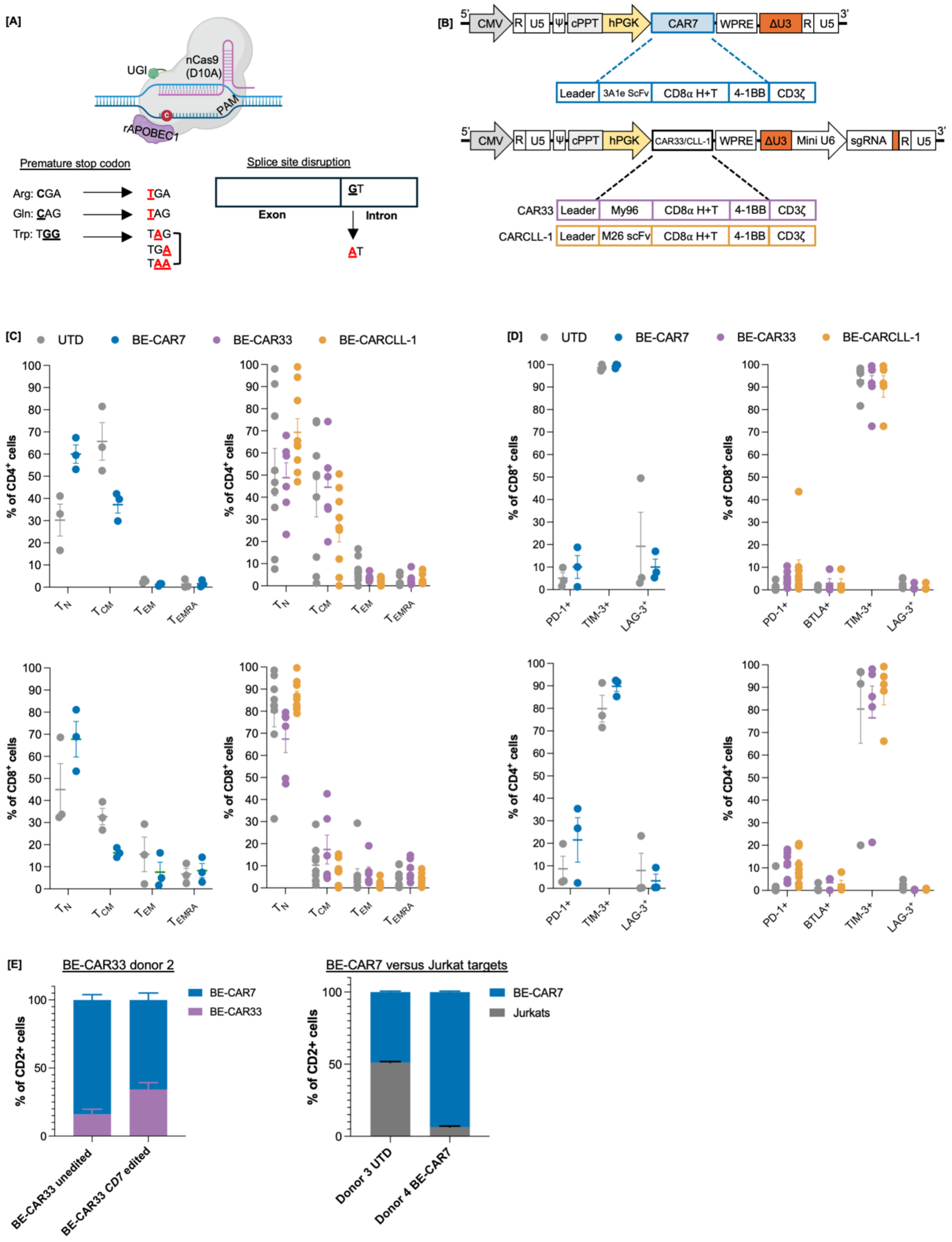

**Supplementary Fig. 1**

**[A]** Strategies employed to cause protein knockout using a cytidine base editor, through the C-to-T mediated conversion of arginine (arg), glutamine (gln) or tryptophan (trp) amino acid residues to premature stop codons (TGG, TGA, TAG), or the disruption of consensus sequences required for proper mRNA splicing. **[B]** Self-inactivating lentiviral vector transfer plasmids utilised for the manufacture of BE-CAR7 (top) and BE-CAR33 or BE-CARCLL-1 (bottom) T cells. The CAR-encoding gene was placed under the control of an internal human phosphoglycerate kinase (hPGK) RNA polymerase II promoter. For pTTB-CAR33 and pTTB-CARCLL-1 plasmids, a sgRNA expression cassette against *TRBC* (with a modified c+5 scaffold) was embedded within the ΔU3 sequence of the 3' LTR under the control of an internal "mini U6" RNA polymerase III promoter (referred to as CRISPR CAR). CMV; cytomegalovirus. **[C]** $CD4^+$ and $CD8^+$ donor-matched untransduced cells (n=3, n=9), BE-CAR7 (n=3) (left), and BE-CAR33 (n=6) as well as BE-CARCLL-1 (n=9) cells (right) displaying combinations of CD62L and/or CD45RA expression profiles associated with $T_N$, $T_{CM}$, $T_{EM}$ or $T_{EMRA}$ subsets. **[D]** $CD4^+$ (top) and $CD8^+$ (bottom) donor-matched untransduced cells, BE-CAR7 (left), as well BE-CAR33 and BE-CARCLL-1 cells (right) displaying cell surface PD-1 (n=3, n=8, n=9 across respective BE-CAR products), BTLA (n=4 across BE-CAR33 and BE-CARCLL-1 only), TIM-3 (n=3, n=5, n=5 across respective BE-CAR products), and LAG-3 (n=3, n=5, n=5 across respective BE-CAR products). **[E] (**Left) CFSE labelled BE-CAR33 targets (T), unedited or edited to disrupt CD7 expression, and BE-CAR7 effectors (E) remaining after co-culture for 16 hours at an E:T of 1:1. Data is shown as technical triplicates from one BE-CAR33 and one BE-CAR7 donor. (Right) $GFP^+$ Jurkat targets (T) remaining after co-culture with unlabelled UTD of BE-CAR7 cells (E) at an E:T of 1:1 for 16 hours. Data is shown as technical triplicates from one UTD and one BE-CAR7 donor.

**Supplemental figure 2**

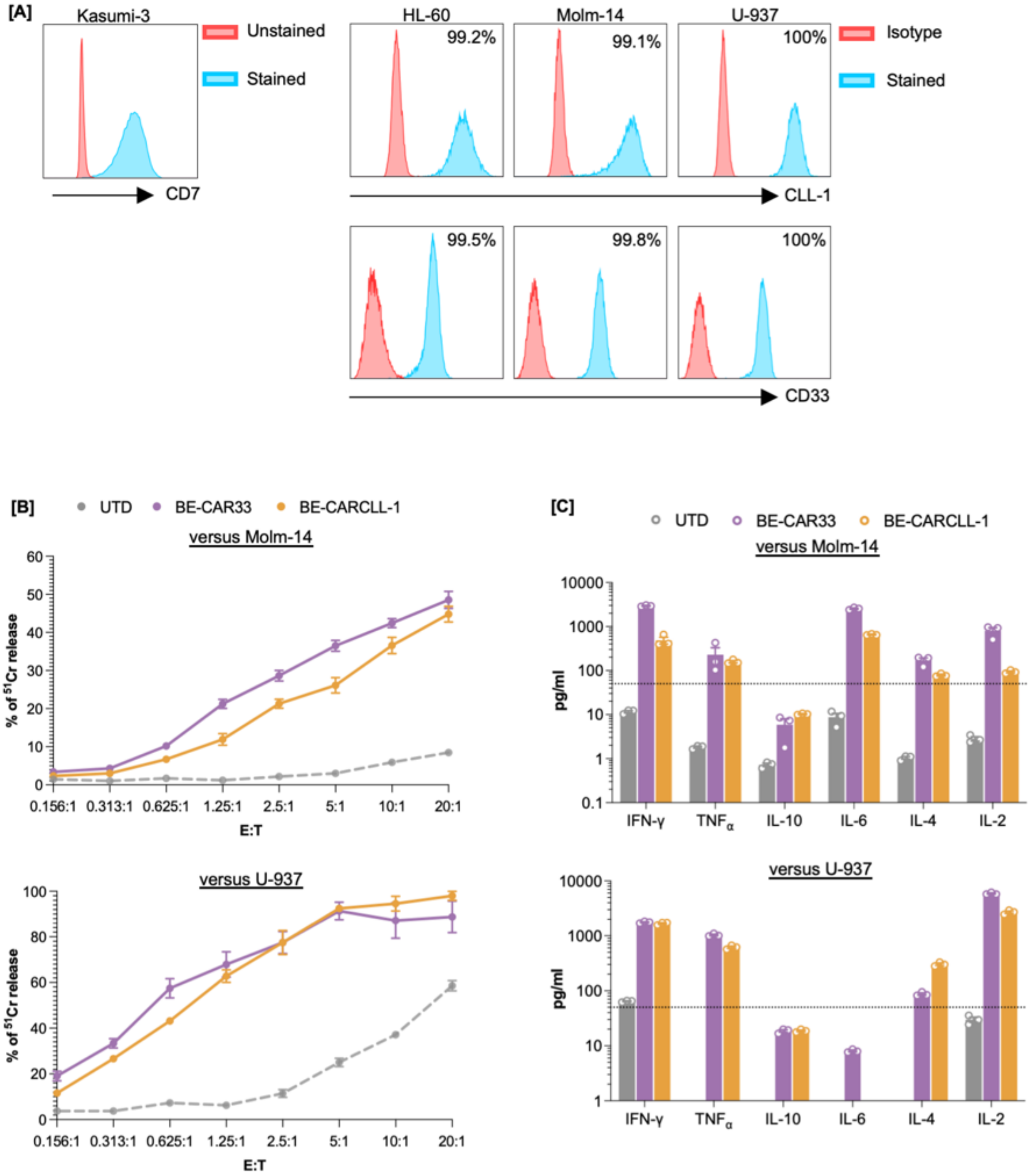

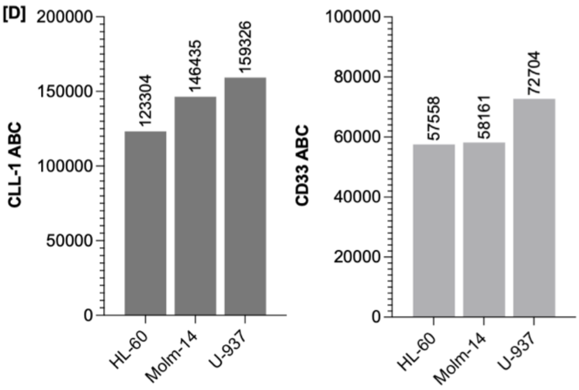


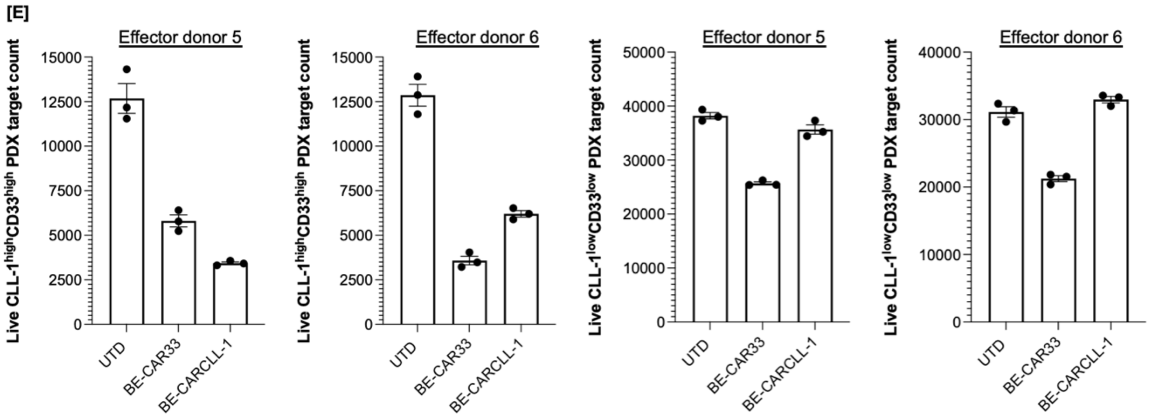


**Supplementary Fig. 2 - legend**

**[A]** Kasumi-3 AML cells demonstrating cell surface CD7, detected using the CD7-6B7 antibody clone, as well as HL-60, Molm-14 and U-937 AML cell lines demonstrating cell surface CLL-1 and CD33 expression detected using 50C1 or P67.6 antibody clones respectively. Analysis of HL-60, Molm-14 and U-937 cells was normalised using matched antibody isotype control. **[B]** Normalised $^{51}$Cr release from pre-labelled Molm-14 (top) or U-937 (bottom) targets when co-cultured with BE-CAR33 or BE-CARCLL-1 for 4 hours at E:Ts ranging from 20:1 to 0.156:1. Untransduced cells were included to demonstrate target lysis in the absence of a CAR. **[C]** Normalised number of (CD45$^{dim}$) CLL-1$^{+}$CD33$^{+}$ or CLL-1$^{dim}$CD33$^{dim}$ PDX target cells remaining following a 16-hour co-culture with untransduced cells, BE-CLL-1CAR or BE-CD33CAR T cells at an E:T of 10:1. Data is shown from two donors with technical triplicates. **[D]** Antibody binding capacity (ABC) of the 50C1 anti-CLL-1 antibody clone (left) and P67.6 anti-CD33 antibody clone (right) on HL-60, Molm-14 and U-937 AML cell lines.

**Supplemental figure 3**

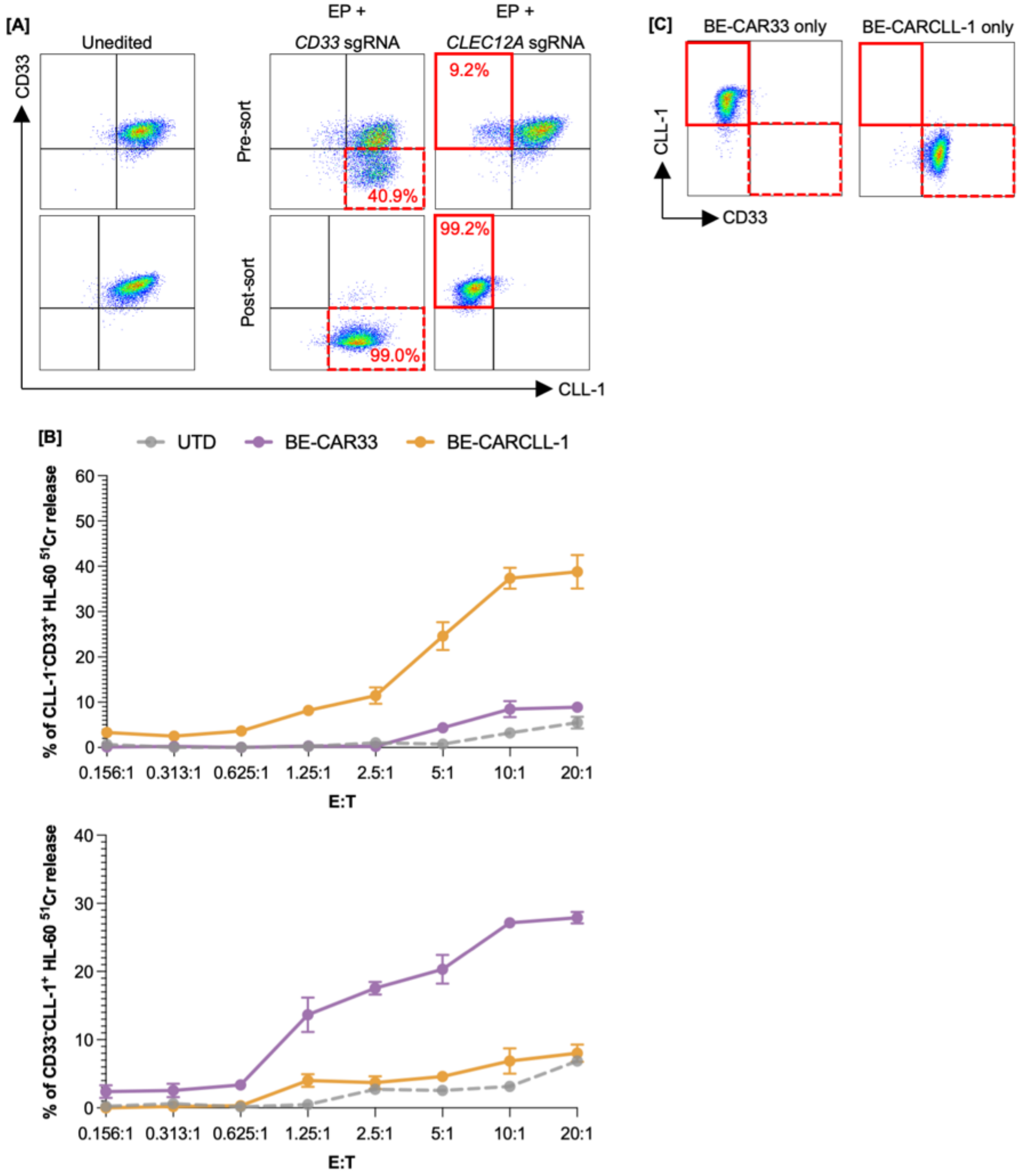


**Supplementary Fig. 3 - legend**

**[A]** (Top) HL-60 cells displaying cell surface expression of CLL-1 and/or CD33 4 days post electroporation with mRNA encoding codon-optimised SpCas9 or BE3, and sgRNA sequences against *CLEC12A* or *CD33*, respectively. Populations harbouring CLL-1$^{-}$CD33$^{+}$ and CD33$^{-}$ CLL-1$^{+}$ phenotypes are highlighted in solid and dashed boxes respectively. (Bottom) HL-60 cells displaying a CLL-1$^-$CD33$^+$ (solid red) or CD33$^-$CLL-1$^+$ (dashed red) phenotype following FACS-assisted enrichment for respective phenotypes. **[B]** Normalised $^{51}$Cr release from pre-labelled CLL-1$^-$CD33$^+$ (top) and CD33$^-$CLL-1$^+$ (bottom) HL-60 targets when co-cultured with BE-CARCLL-1 or BE-CAR33 T cells for 4 hours at E:Ts ranging from 20:1 to 0.156:1. Untransduced cells were included to demonstrate target lysis in the absence of a CAR. **[C]** (Live CD45$^{dim/+}$) GFP$^+$ tumour detected in bone marrows of mice treated with BE-CARCLL- 1 or BE-CAR33 T cells in isolation displaying cell surface CLL-1 or CD33 expression.
